# An automated platform for spatial functional modeling and fingerprint analysis of tissue molecular landscapes

**DOI:** 10.64898/2026.07.15.738836

**Authors:** Morteza Hajihosseini, Eduardo Patino-Martinez, Rahul Ghosal, Mariana J. Kaplan, Saumyadipta Pyne

## Abstract

Spatial transcriptomics (ST) enables high-resolution molecular profiling while preserving tissue architecture, creating new opportunities to investigate how disease-associated pathways are organized within tissues. However, existing analytical approaches largely focus on individual pathways or cell types and do not provide a unified framework for modeling spatially varying pathway interactions across tissue sections and anatomical planes. We present an integrative framework, Spatial Fingerprints Analytics (SFinx), that introduces the concept of a spatial fingerprint for representing patterns of molecular signatures, and Spatial Functional Data Analysis for spatial regression and mapping of localized pathway activity and pathway–phenotype interactions in complex tissues. Applying SFinx to ST datasets on murine lupus nephritis, we reconstructed continuous spatial landscapes of pathway activity and disease-associated phenotypes across kidney sections. This approach identified anatomically restricted inflammatory domains characterized by coordinated activation of immune pathways and revealed substantial spatial heterogeneity in pathway crosstalk across renal compartments. Using generalized additive models with tensor-product splines, we quantified spatially varying associations between lupus nephritis and neutrophil activation pathways across tissue sections, uncovering regions with both positive and negative relationships that would be obscured by conventional bulk analyses. Multi-slice integration further demonstrated reproducible spatial interaction patterns while accounting for section-specific variability. Together, SFinx transforms mixed-spot transcriptomic measurements into interpretable spatial pathway landscapes and interaction maps, providing a general framework for identifying localized disease mechanisms. SFinx revealed previously unrecognized spatial organization of inflammatory signaling in lupus nephritis and presents a broadly applicable strategy for studying spatially coordinated biological processes in cancers, autoimmune and neurodegenerative diseases.

## 1. Introduction

Recent advances in spatial transcriptomics (ST) have transformed tissue-scale molecular profiling by enabling transcriptome-wide measurements while preserving spatial context [1]. Platforms such as 10x Visium, Slide-seq, and MERFISH now allow mapping of molecular programs across intact tissues at resolutions ranging from multicellular regions to near single-cell levels, revealing cellular neighborhoods, spatial gradients, and microenvironmental interactions that were previously inaccessible [1,2]. ST has also enabled detailed characterization of tumor microenvironments by uncovering complex spatial heterogeneity in molecular programs, immune cell infiltration, and cell–cell interactions, providing important insights into tumor progression, metastatic potential, and therapeutic response across multiple cancer types [3].

Despite these advances, extracting biologically meaningful information from ST data remains challenging. Spatially adjacent locations exhibit intrinsic autocorrelation, measurements often represent mixtures of cell states, and disease-relevant processes frequently emerge at the level of pathways and tissue niches rather than individual genes. Existing methods address important components of ST analysis. SpaGCN identifies spatial domains and spatially variable genes [1], Giotto provides a comprehensive analysis environment [4], and stLearn integrates morphology, trajectories, and cell–cell interactions [5]. However, these approaches do not directly model spatially varying pathway–pathway or pathway–phenotype relationships across tissue space.

We previously developed geostatistical approaches for modeling continuous molecular landscapes and quantifying spatial heterogeneity within tissues [6,7]. Although these methods revealed biologically meaningful spatial structure beyond gene-level analyses, they were not designed to characterize pathway activity or interactions between pathways across tissue space. In parallel, our recent work on longitudinal multi-omic and spatial transcriptomic study of lupus nephritis has revealed enrichment of inflammatory pathways and spatially organized immune niches in this condition [8]. However, existing analytical tools are limited in their ability to systematically characterize how disease-relevant pathways interact across anatomical regions, particularly when these relationships vary spatially.

In this study, we introduce a statistical framework based on two novel features. First, the Spatial Fingerprints Analytics (SFinx) framework introduces the concept of a *spatial fingerprint*, a quantitative representation of a spatial pattern of the enrichment of a molecular signature across the tissue space. Indeed, for the given transcriptome data of a tissue slice under study and a known molecular signature *S*, SFinx first discretizes the tissue space into spatial units with well-defined location and shape each of which summarizes the statistical significance of the local expression of *S*. Together, the matrix of spatial significance scores induces a programmatically generated fingerprint (*FP_S_*) of *S* on the tissue space. Importantly, such fingerprints involving one or more signatures allow systematic application of logical and/or statistical operators (defined on the underlying tissue space) thereby recursively defining a complex pattern with a precise, objective statement to describe a potentially subjective visual phenomenon. Examples of such operators include both unary (complement, autocorrelation) and binary (union, intersection, difference, maximum, etc.). This makes it possible to textually encode a spatial hypothesis for the purpose of statistical testing or a complex visual pattern for quantitative investigation. Notably, a fingerprint generated by SFinx offers a programmable (with flexible choices of statistical thresholds and logical operators) multimodal (dual text-image) representation of a target molecular landscape.

A second key innovation is Spatial Functional Data Analysis (SFDA), which represents pathway activity as continuous two-dimensional surfaces and enables statistical modeling of spatially varying pathway relationships. SFinx implements the SFDA framework, built on the capability of *functional statistics*, an established field well-known for its rigorous representation and analysis of data as curves and surfaces [9]. Thus, SFDA enables SFinx to quantify and visualize the heterogeneity on a molecular landscape in a given tissue space, as well as to characterize spatially varying associations between biological pathways across tissue slices. Few applications exist today for handling contiguous 2D slices of a 3D tissue sample under investigation, thus enabling reconstruction of the overall molecular landscape including inter-slice information along the “z-axis”. To integrate information across tissue slices, current analytical approaches either focus on their computational alignment [10,11] or use the average expression values among them. Such inter-slice variation in expression should ideally be accounted for systematically with the help of slice-specific terms as used in our spatial functional regression model.

Interestingly, the capability of functionally representing a molecular landscape of a given gene signature not only allows SFinx to analyze the spatial expression of a selected pathway (as well as the co-expression of multiple pathways), but it also simultaneously enables the mapping of spatial phenotypes on the same tissue space. For instance, our platform leverages an established and curated database named ‘Human Phenotype Ontology’ (HPO) for its annotations of hundreds of known clinical phenotypes in terms of the gene signatures that are associated with each of them [12]. This leads to automatic application of functional regression by SFinx to test for association between the landscapes of one or more disease pathways as predictors of a tissue phenotype observed as the spatial outcome.

We applied SFinx to lupus nephritis, a prototypic autoimmune disease characterized by spatially heterogeneous immune infiltration and tissue injury, providing an ideal system for evaluating localized pathway crosstalk [13]. Known phenotypes include deposition of autoantibodies and immune complexes, cytokine production, activation and proliferation of infiltrating immune cells that amplify and sustain inflammation. Our approach enables the direct mapping of inflammatory and immune-regulatory pathways onto histological structures, thereby facilitating the identification of spatially localized crosstalk between disease-relevant processes, including neutrophil activation and autoimmune-mediated tissue injury.

Using 10x Visium data generated from a murine lupus nephritis model, we sought to (i) reconstruct continuous pathway activity landscapes, (ii) identify spatial domains characterized by convergent or divergent inflammatory signaling, and (iii) quantify spatially varying relationships between neutrophil activation and lupus-associated disease phenotypes using SFDA. Together, SFinx extends spatial transcriptomic analysis from descriptive mapping of genes and cell types to quantitative modeling of spatial pathway interactions and disease-associated molecular landscapes. This framework provides a general strategy for identifying localized disease mechanisms in autoimmune disorders, cancer, neurodegeneration, and other spatially organized pathologies.

## 2. Materials and methods

### 2.1. Data

We analyzed 10X Visium ST data across four consecutive tissue slices of a kidney sample from a LN mouse model denoted in this study by slices 1, 2, 3 and 4. Each slice had spatial data available at 4726, 4760, 4637, and 4637 locations respectively. These were preprocessed through STdeconvolve to identify cell-type composition and reconstruct spot-level gene expression profiles. For each slice, this step yielded data for the top 5000 over-dispersed genes. While the gene expression values were averaged over the slices in our previous analysis, here we studied them using a slice-specific model. More details of tissue preparation and ST data generation and preprocessing are available in our previous publication [8]. In brief, female lupus-prone NZB/W F1 mice (The Jackson Laboratory, Bar Harbor, ME, USA) were maintained under specific pathogen-free conditions at the NIH animal facility. All procedures were approved by the NIAMS Animal Care and Use Committee (Protocol No. A019-05-03). Mice were euthanized at 10, 20, and 30 weeks of age, and blood and tissue samples were collected for analysis. Kidneys were embedded in Optimal Cutting Temperature (OCT) compound, cryosectioned (15 µm), and mounted onto Visium Spatial Gene Expression slides. Sections were methanol-fixed, hematoxylin and eosin (H&E)-stained, imaged, and processed for spatial transcriptomics following the manufacturer’s protocol. cDNA libraries were sequenced on an Illumina NovaSeq 6000, and gene expression data were visualized using Loupe Browser. The steps of data collection are summarized in **Figure 1**.

**Figure 1.**
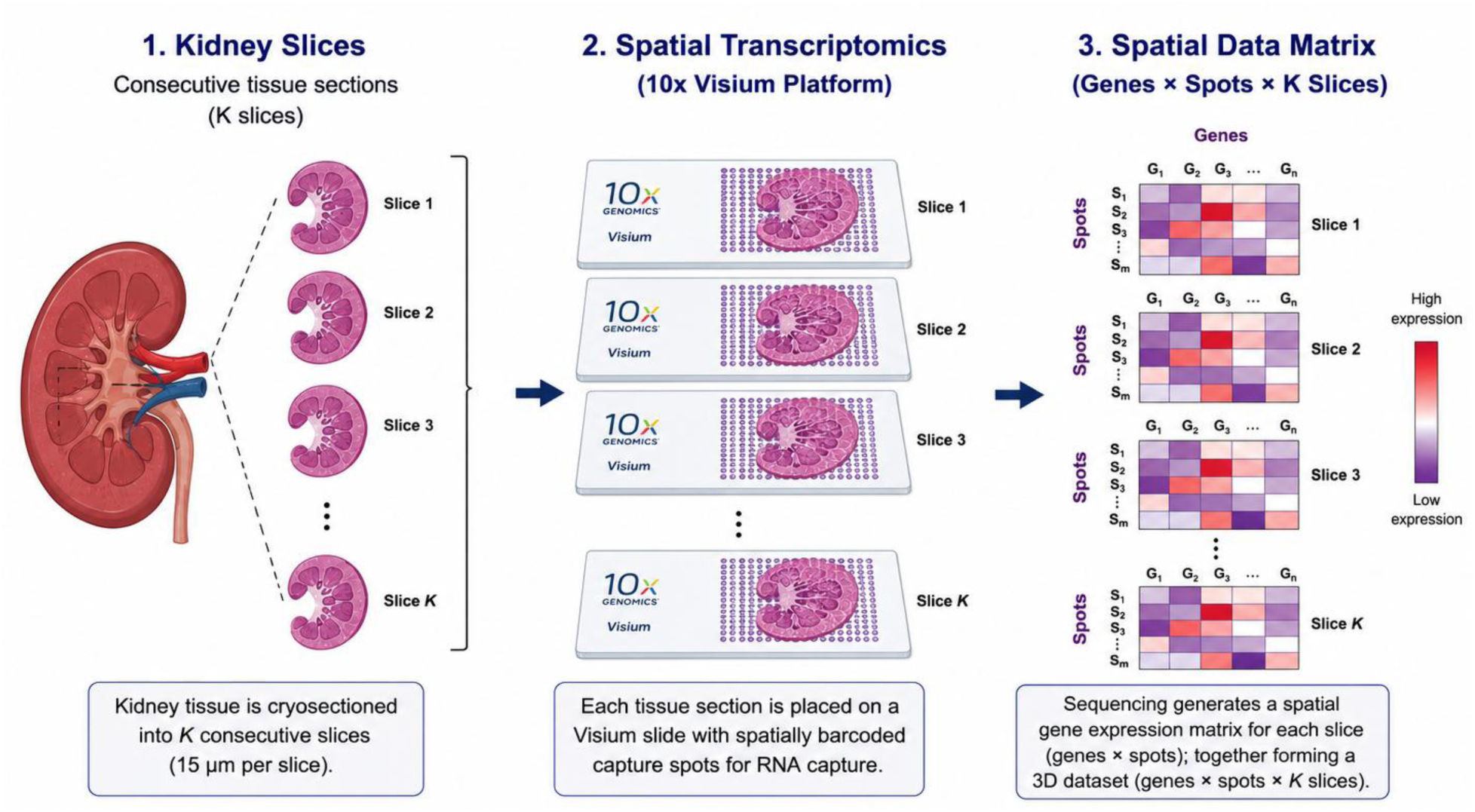
Spatial transcriptomics workflow for kidney tissue analysis. Kidney tissue was collected from a lupus-prone mouse model and serially sectioned into consecutive slices. Tissue sections were processed using the 10x Genomics Visium spatial transcriptomics platform, where mRNA molecules were captured at spatially barcoded spots while preserving tissue architecture. Sequencing generated a spatial gene expression matrix consisting of genes (columns) and spatial capture spots (rows), providing the foundation for downstream spatial fingerprinting, pathway enrichment analysis, and spatial functional data analysis.

### 2.2. Spatial Fingerprints (SFinx)

The SFinx pipeline consists of four major components. First, we employ STdeconvolve, a reference-free approach to deconvolve underlying cell types and their transcriptomic profiles from multi-cellular spatial transcriptomics data [14]. Second, we reconstruct spot-level gene expression by computing the matrix product of cell-type proportions and cell-type-specific gene expression profiles. Third, we apply ordered quantile (ORQ) normalization [15] followed by z-score transformation to ensure comparability across spatial locations and tissue sections. Fourth, we compute single-sample pathway enrichment scores using singscore algorithm [16,17], a rank-based metric of gene set enrichment that scores the expression activities of gene sets at a single-sample level, enabling us to quantify pathway activity at each spatial location independent of other samples.

Following pathway scoring, we apply ordinary kriging interpolation with distance-bounded grids to generate smooth, continuous pathway activity surfaces [18]. This approach creates prediction grids that only include points within a specified distance threshold from actual tissue spots, preventing extrapolation into non-tissue regions. The distance threshold is considered as 1.2 times the average nearest-neighbor distance, ensuring biological relevance of interpolated values while enabling integration of multiple tissue sections onto common spatial coordinates.

SFinx also implements a suite of binary operators for pathway fingerprint analysis. For two pathway fingerprints defined by threshold-based activation (|*Z*| ≥ threshold), we computed: (1) Union: locations where either pathway is activated; (2) Intersection: locations where both pathways co-activate, quantified by Jaccard similarity index[15]; (3) Difference: locations with asymmetric activation; and (4) Maximum: the dominant pathway at each location.

To characterize the sensitivity of spatial fingerprints to threshold selection, we additionally visualized each pathway fingerprint for one of the slices at three rigour levels including low (|*Z*| ≥ 0.5), medium (|*Z*| ≥ 1.0), and high (|*Z*| ≥ 1.5) using two-dimensional kernel density estimates of retained spots using plasma color scale, enabling direct visual assessment of how activation threshold affects the spatial extent and localization of each pathway fingerprint.

### 2.3. Spatial Functional Data Analysis (SFDA)

The core innovation of our pipeline lies in its application of spatial functional data analysis (SFDA) [19,20] which uses spatially varying coefficients to model the pathway-pathway relationships. Let us denote the spatially varying response pathway enrichment score for a particular slice as *Y*(*s*_1,_*s*_2_) and the predictor pathway score as *X*(*s*_1,_*s*_2_) at longitude *s*_1,_ and latitude *s*_2_. We use the following spatially varying function-on-function regression model [19] to quantify the spatially varying association between the response and predictor pathway scores:

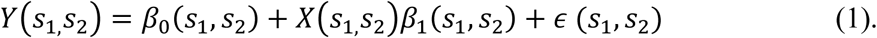

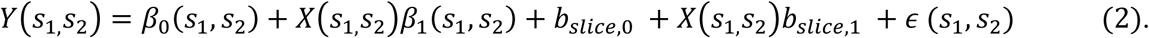

In the above models *β*_1_(*s*_1_, *s*_2_) is a two-dimensional functional slope which captures the spatially varying associating between the response and the predictor pathway enrichment score, *β*_0_(*s*_1_, *s*_2_) is a functional intercept, and *ε* (*s*_1_, *s*_2_) is a mean zero measurement error. The slice-specific random intercept and slope is denoted by *b_slice_*_,0_ and *b_slice_*_,1_. The above models can be seen as extension of Generalized additive models (GAMs) [21,22], which are flexible semiparametric methods that allow the inclusion of both fixed and spatially varying coefficients. The spatially varying coefficients are modelled using tensor product splines and the coefficients can be estimated using a penalized least square approach, with explicit smoothness penalty which allows the coefficients to vary smoothly over space [19,23], through smoothing functions such as splines [24]. The above model can be efficiently implemented using the *bam* function within the *mgcv* package [25] in *R*, where the response pathway expression is modeled as a function of spatial coordinates (longitude and latitude) and their interaction with predictor pathway expression. We used 20 cubic B-spline basis functions in both spatial dimensions and chose the smoothing parameters using the REML method [25].

Specifically, the model command in gam for equation [2] takes the form:

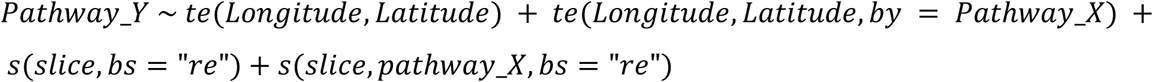

where pathway_Y represents the response pathway enrichment score, pathway_X represents the predictor pathway score, and *te*() command specifies the tensor product smoothing spline estimation for the bivariate coefficients. This formulation allows the relationship between predictor and response pathways to vary smoothly across tissue space, capturing regional differences in pathway crosstalk that may reflect distinct disease microenvironments or cellular states. The spatially varying coefficient surfaces are extracted through model on a 50×50 spatial grid, with statistical significance assessed through point-wise 95% confidence intervals [26].

Regions where confidence intervals exclude zero indicate spatial locations with significant pathway-pathway associations. We computed significance surfaces as binary masks that identify locations with robust pointwise signals and further classified these regions into areas exhibiting positive and negative significant effects. This approach reveals not just whether pathways interact, but precisely where in the tissue these interactions occur and how their directionality varies across anatomical compartments.

All statistical analyses and visualizations were performed in R version 4.5.2 (R Foundation for Statistical Computing, Vienna, Austria). R packages used in the analytical workflow included *singscore* for gene signature scoring, *mgcv* for generalized additive modeling and spatially varying coefficient estimation, *automap* and *gstat* for spatial kriging interpolation, *sf* and *sp* for spatial data handling, *ggplot2* and *plot3D* for data visualization.

## 3. Results

To demonstrate the utility of SFinx and SFDA, we applied our framework to spatial transcriptomic data generated from four kidney sections of a murine lupus nephritis (LN) model [8]. LN is one of the most severe manifestations of systemic lupus erythematosus and is characterized by spatially heterogeneous immune infiltration, inflammatory signaling, and tissue injury, making it an ideal system for evaluating localized pathway interactions [27].

We focused on two biologically relevant signatures representing disease pathology and inflammatory effector mechanisms. The first, HP_LUPUS_NEPHRITIS, was obtained from the Human Phenotype Ontology [28] comprises genes associated with lupus nephritis pathology. The second, GOBP_REGULATION_OF_NEUTROPHIL_ACTIVATION, was derived from Gene Ontology Biological Processes and captures neutrophil-associated inflammatory pathways implicated in renal injury. Neutrophils accumulate within inflamed kidneys during LN and contribute to tissue damage through release of reactive oxygen species, proteolytic enzymes, and other inflammatory mediators [29]. Together, these signatures enabled us to investigate how disease-associated molecular programs are spatially organized and how neutrophil-driven inflammation relates to localized LN pathology.

For each tissue section, pathway activity was quantified at every spatial location using single-sample enrichment analysis (singscore) applied to rank-transformed gene expression profiles. These pathway activity estimates were subsequently projected onto a common spatial grid and used as inputs for SFinx fingerprint generation and downstream SFDA analyses. To evaluate how threshold strictness influences spatial fingerprint, we examined FP1 (Mouse Neutrophil Activation Pathway) and FP2 (Lupus Nephritis Pathway) in kidney Slice 3 across three activation thresholds: Low (|*Z*| ≥ 0.5), Medium (|*Z*| ≥ 1.0), and High (|*Z*| ≥ 1.5) in **Figure 2**.

**Figure 2.**
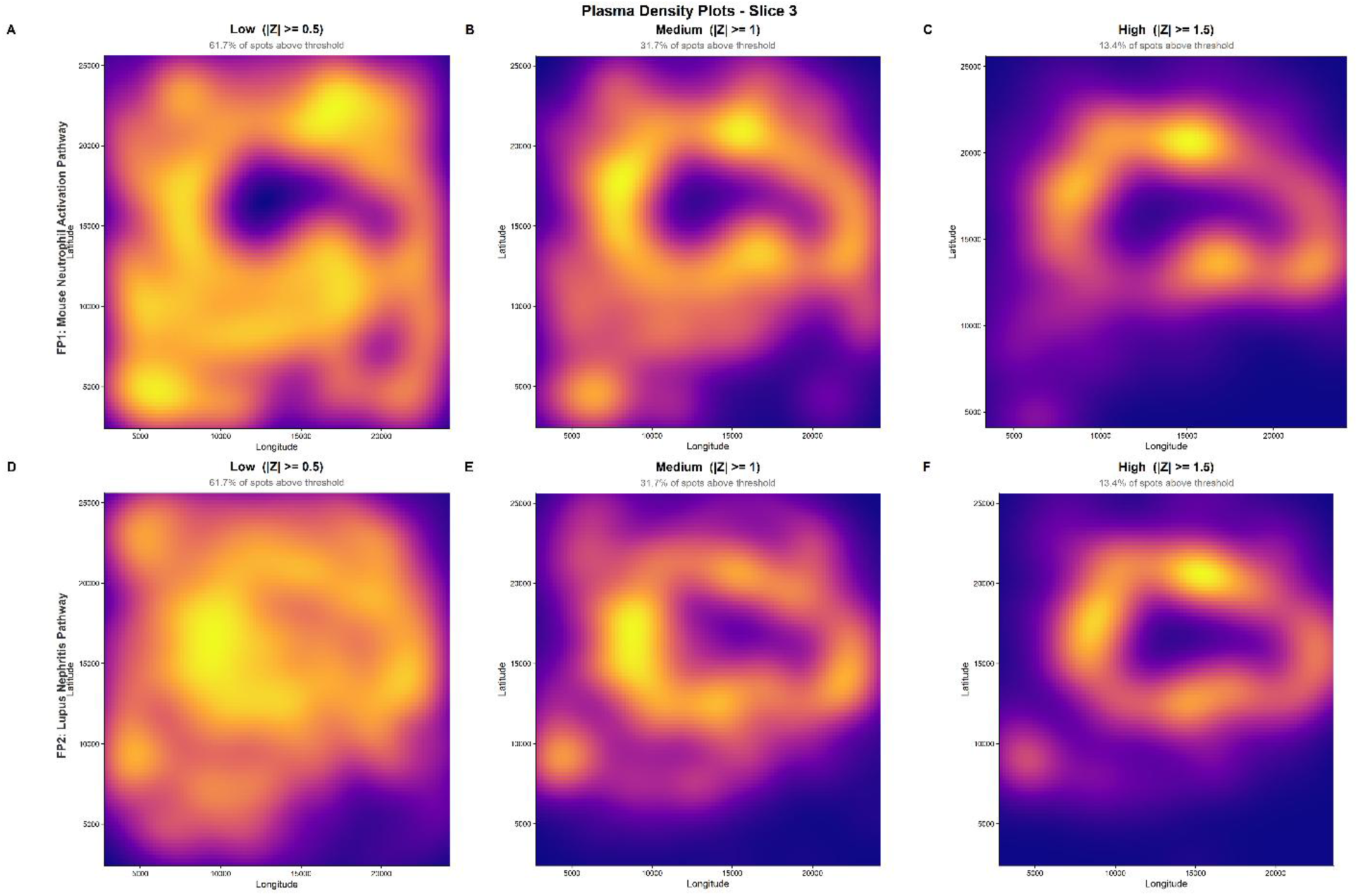
Plasma density maps of two fingerprint pathways in Slice 3 at increasing activation thresholds. Each panel shows a 2D kernel density estimate of spatially resolved spots exceeding an absolute Z-score threshold (|*Z*| ≥ 0.5, 1.0, and 1.5 for Low, Medium, and High, respectively) for FP1: Mouse Neutrophil Activation Pathway (top row) and FP2: Lupus Nephritis Pathway (bottom row). Color intensity reflects local spot density, ranging from low (dark purple) to high (bright yellow), using the plasma color scale. The percentage of spots meeting each threshold is indicated in the subtitle of each panel. As the threshold increases, fewer spots are retained, revealing the spatial fingerprint of strongest pathway activation within the tissue section.

At the lowest threshold, 61.7% of spots were retained for both pathways, revealing broad spatial coverage of the tissue section. As strictness increased to Medium and High thresholds, the proportion of retained spots fell to 31.7% and 13.4% respectively and progressively isolating the spatial hotspots of strongest pathway activation. For FP1, high-threshold panels revealed concentrated neutrophil activation in discrete anatomical zones, consistent with localized inflammatory niches. For FP2, the lupus nephritis signature reduced at high thresholds into spatially distinct clusters that do not fully overlap with FP1, suggesting at least partially non-overlapping spatial domains of pathway activity. These observations demonstrate that spatial fingerprints capture pathway-specific spatial organization and that threshold selection can be used to interrogate biological processes across multiple spatial scales. The distinct spatial patterns observed for FP1 and FP2 motivated subsequent analyses using SFinx unary and binary operators to systematically characterize pathway activation, overlap, and spatial exclusion. Unary operator results for all pathway–slice combinations at the medium threshold (|*Z*| ≥ 1.0) are shown in **Figure 3**.

**Figure 3:**
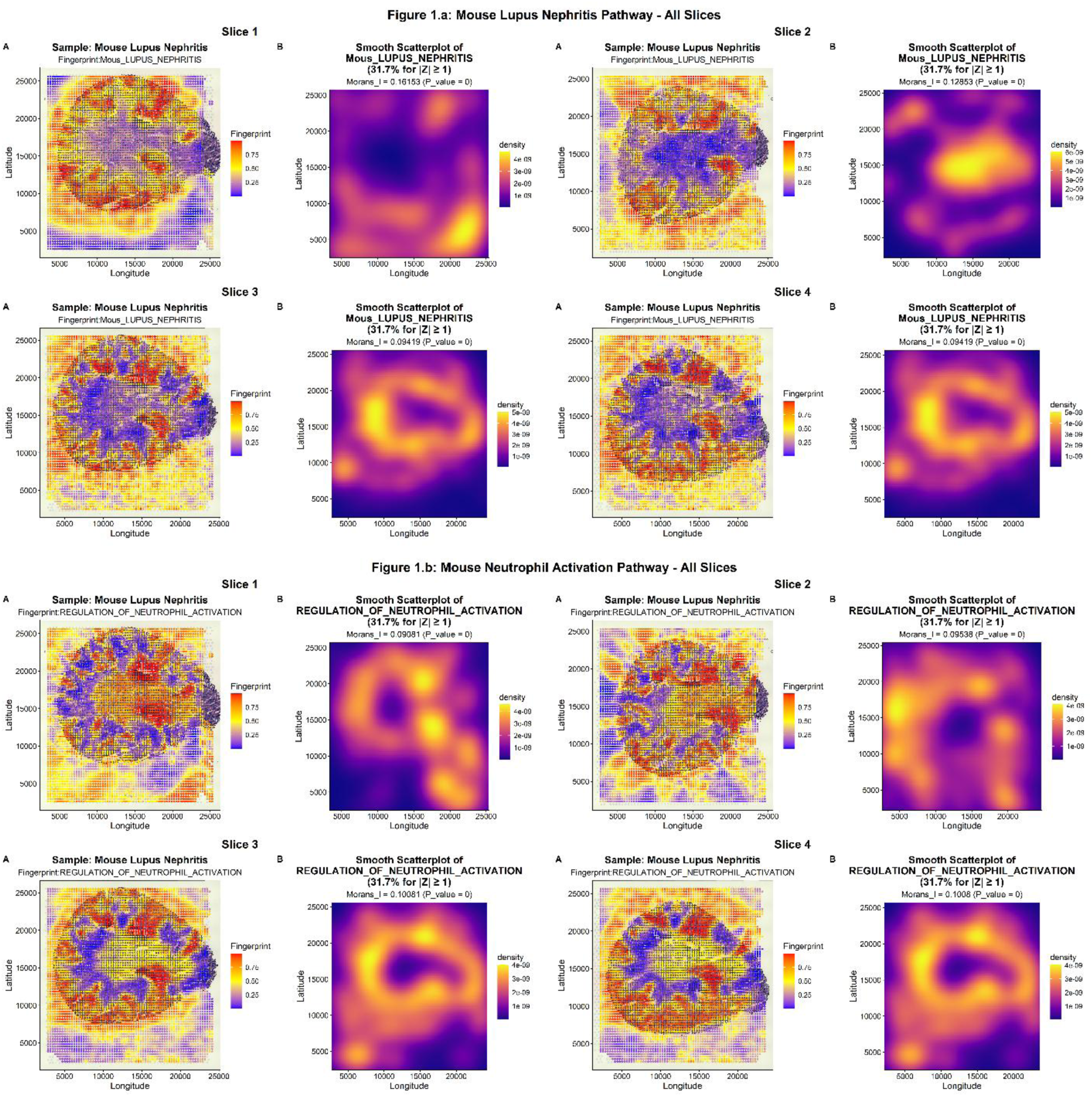
SFinx unary operators’ analysis for Mouse Lupus Nephritis Pathway and Mouse Neutrophil Activation Pathway in each slice. **Panel A**: Spatial distribution of pathway enrichment scores across each kidney tissue section. Colors represent pathway activity scores transformed to probability scale (blue = low activity, yellow = medium, red = high activity). Points indicate individual spatial transcriptomics spots overlaid on H&E-stained tissue histology. **Panel B**: Density heatmap of activated regions for pathway enrichment scores (|Z| ≥ 1.0). Plasma color scale indicates spatial concentration of pathway activation, revealing percentage of tissue spots exceed the activation threshold. The Moran’s I value indicates spatial clustering of pathway activity.

For each tissue slice in **Figure 3**, unary operator analysis revealed the spatial distribution and clustering patterns of individual pathway activities. Pathway fingerprints were defined as locations with pathway scores |Z| ≥ 1, representing regions of significant pathway activation. Moran’s *I* statistic quantified spatial autocorrelation, confirming non-random clustering of pathway activity. Smooth scatterplots and contour maps identified anatomical hotspots of lupus nephritis signatures and neutrophil activation, with density visualization revealing focal inflammatory zones. Figure 4 explored more pathway-pathway interactions by using SFinx binary operators including union, intersection, difference, and maximum of spatial domains of pathway activities.

**Figure 4:**
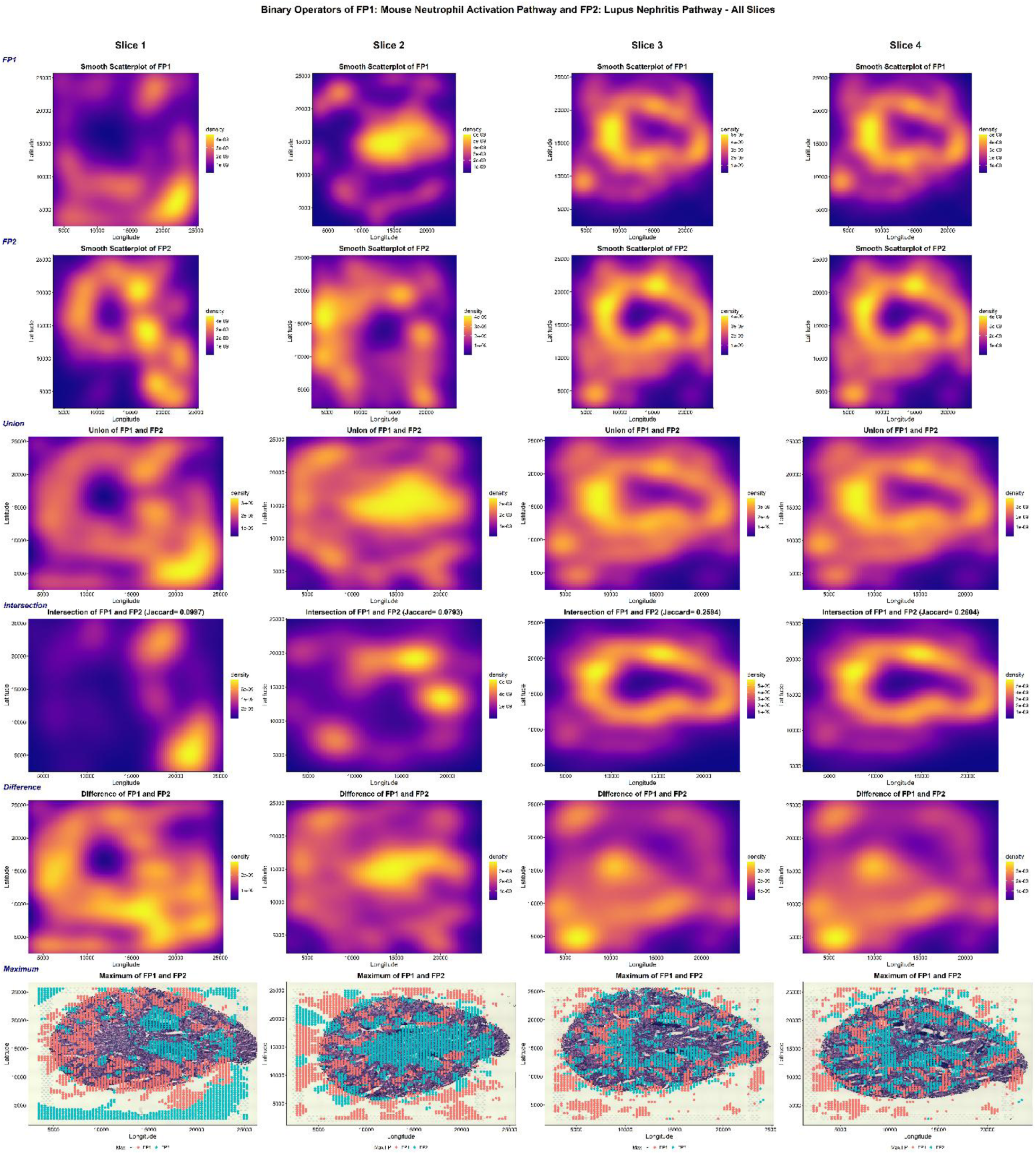
SFinx binary operators’ analysis for each slice. **Panel A**: Spatial density distribution of activated regions for fingerprint 1. Plasma color gradient indicates concentration of spots exceeding activation threshold (|*Z*| ≥ 1.0), revealing focal hotspots of fingerprint 1 activity. **Panel B**: Spatial density distribution of activated regions for fingerprint 2. Plasma color gradient indicates concentration of spots exceeding activation threshold (|*Z*| ≥ 1.0), revealing focal hotspots of fingerprint 2 activity. **Panel C**: Union operator showing combined regions where either fingerprint 1 or fingerprint 2 (or both) are activated. It represents the total inflammatory landscape encompassing both pathway activities across kidney tissue. **Panel D**: Intersection operator identifying co-localized regions where both fingerprint 1 and fingerprint 2 are simultaneously activated. Jaccard similarity index quantifies spatial overlap, with focal hotspots indicating sites of convergent pathway activation. **Panel E**: Difference operator highlighting regions with asymmetric pathway activation (either fingerprint 1 or fingerprint 2 activated, but not both). Reveals spatially distinct inflammatory mechanisms operating in different tissue microenvironments. **Panel F**: Maximum operator showing dominant pathway at each activated location. Colors indicate which pathway (FP1: pathway 1 or FP2: pathway 2) exhibits stronger activation, revealing spatial partitioning of pathway dominance across tissue architecture.

Binary operator analysis in **Figure 4** examined the spatial co-localization and interaction between lupus nephritis and neutrophil activation pathways. Chi-square tests revealed significant spatial dependence between the two pathway fingerprints (*P* < 0.05 across sections), indicating non-random co-activation patterns. Jaccard similarity indices quantified the degree of pathway overlap, ranging from 0.07 to 0.2 across sections, suggesting moderate spatial concordance.

Union maps identified broad regions of inflammatory activity, while intersection maps highlighted specific anatomical compartments with simultaneous activation of both pathways, potentially representing sites of active neutrophil-mediated kidney damage. Difference maps revealed regions with pathway-specific activation, suggesting distinct inflammatory mechanisms operating in different tissue microenvironments.

Our spatial function on function regression revealed substantial spatial heterogeneity in the neutrophil activation-lupus nephritis pathway relationship across kidney tissue. For individual tissue sections, slice-specific models demonstrated spatially varying patterns of association. Coefficient surfaces exhibited both positive and negative associations in different anatomical regions, suggesting that neutrophil activation’s relationship with lupus nephritis signatures depends on local tissue microenvironment. Model summaries provided statistical evidence for the spatially varying associations (*P* < 0.001 for tensor product smooths).

The 3D perspective plots in **Figure 5** displays the spatially varying coefficient landscape, with lower and upper 95% confidence surfaces bracketing the estimated coefficient surface. Cross-sectional profiles along longitude and latitude axes revealed directional trends in pathway relationships, with some regions showing strong positive coefficients (indicating neutrophil activation positively predicts lupus nephritis activity) and others showing negative or null associations. Significance maps identified focal regions where confidence intervals excluded zero, representing anatomical locations with statistically robust pathway interactions.

**Figure 5:**
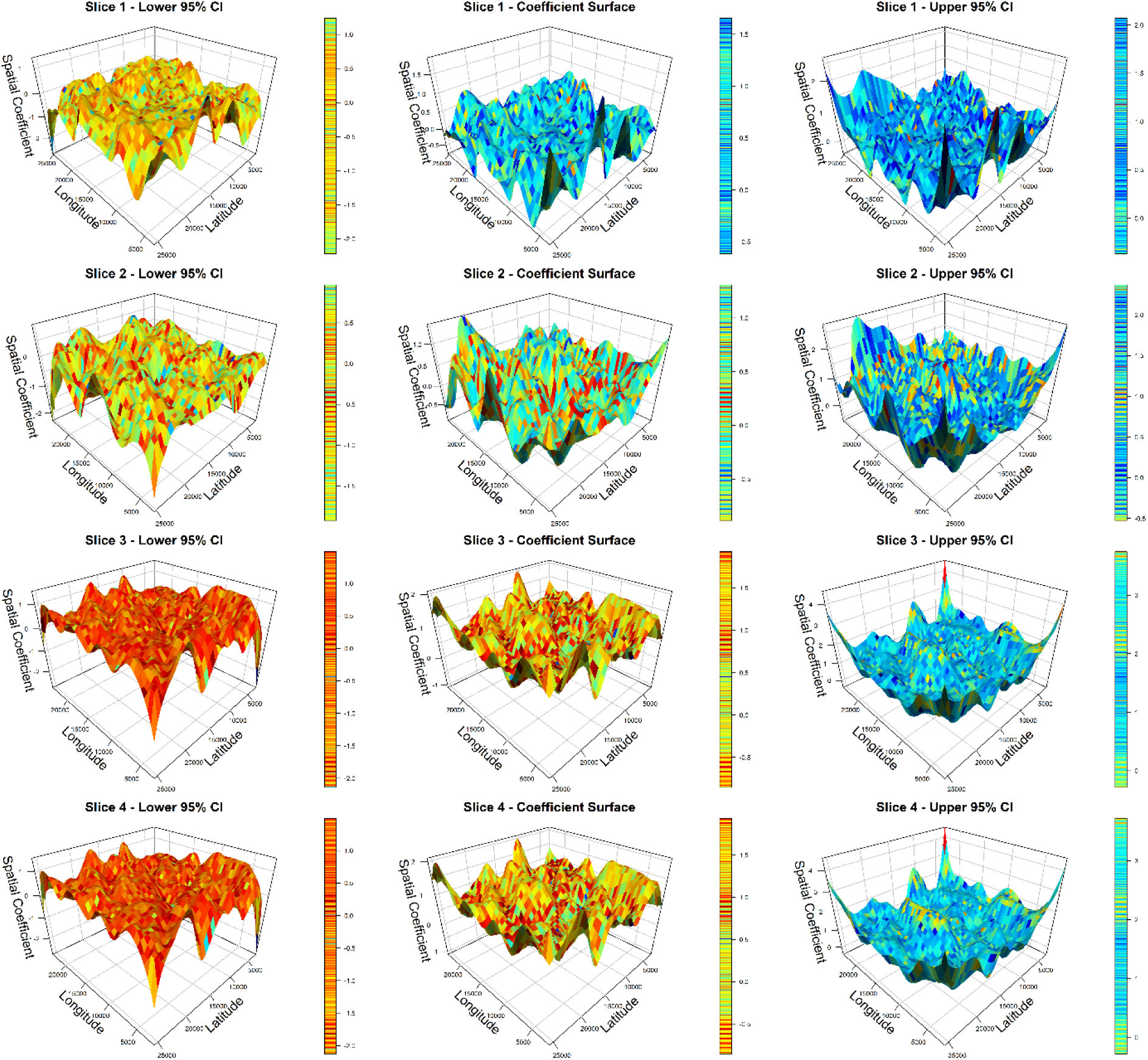
Spatial functional data analysis for each slice. **Left Panel:** Lower bound of 95% confidence interval for spatially varying coefficient surface. It represents conservative estimate of Pathway X’s effect on Pathway Y across kidney tissue geography. **Middle Panel:** Spatially varying coefficient surface quantifying the relationship between Pathway X (predictor) and Pathway Y (response) across tissue space. Positive values (blue peaks) indicate regions where Pathway X activation positively predicts Pathway Y activity; negative values indicate antagonistic relationships. Surface smoothness reflects generalized additive model tensor product splines (*k*=20). **Right Panel:** Upper bound of 95% confidence interval for spatially varying coefficient surface. Regions where lower and upper CI bounds exclude zero indicate statistically significant pathway relationships.

Integration of all four tissue slices in **Figure 6** into a unified multilayer model (with random effects for slice identity and slice-specific pathway slopes) revealed consistent spatial patterns in pathway crosstalk across serial sections. The fixed-effects coefficient surface represented the average spatial relationship between neutrophil activation and lupus nephritis pathways across the tissue volume, while random effects captured section-specific deviations from this average pattern.

**Figure 6:**
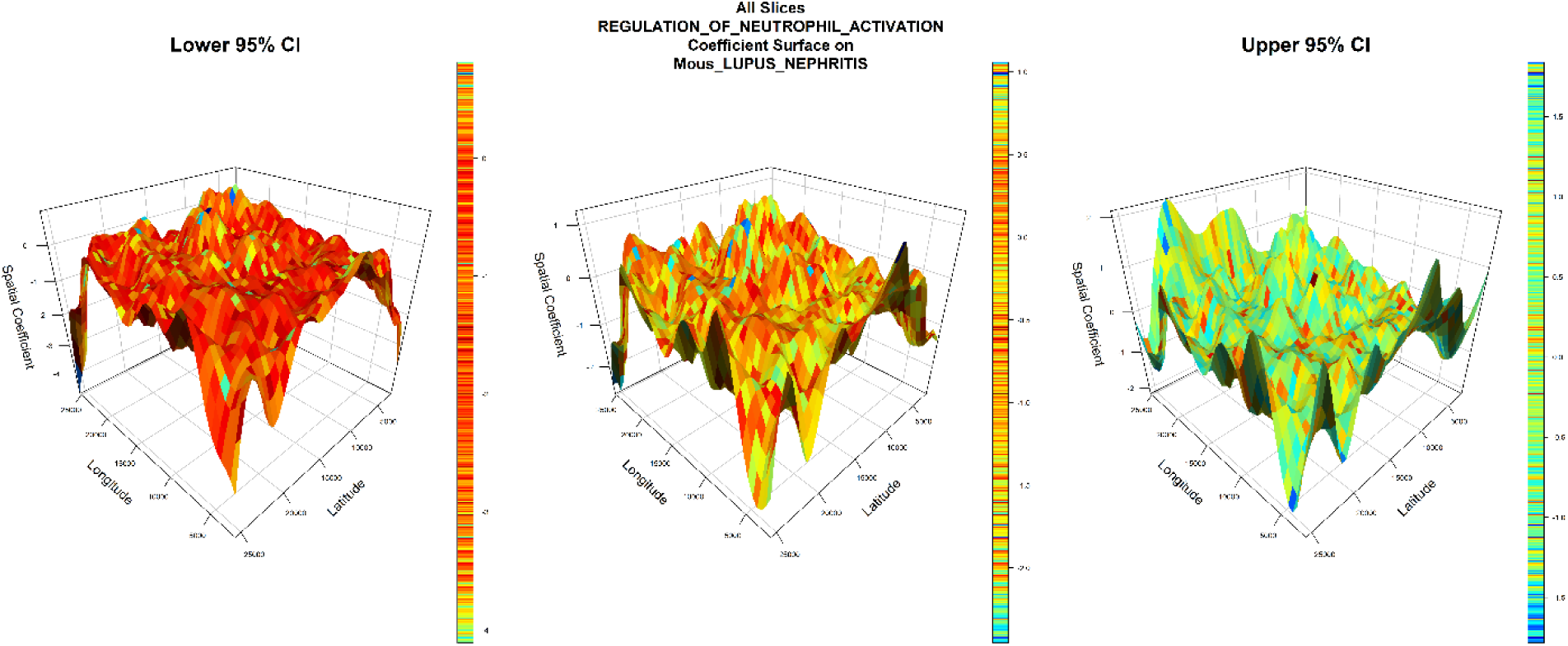
Spatial functional data analysis for all slices with slice-specific random effects. **Left Panel:** Lower bound of 95% confidence interval for spatially varying coefficient surface. Represents conservative estimate of Pathway X effect on Pathway Y across kidney tissue geography. **Middle Panel:** Spatially varying coefficient surface quantifying the relationship between Pathway X (predictor) and Pathway Y (response) across tissue space. Positive values (blue peaks) indicate regions where Pathway X activation positively predicts Pathway Y activity; negative values indicate antagonistic relationships. Surface smoothness reflects generalized additive model tensor product splines (k=20). **Right Panel:** Upper bound of 95% confidence interval for spatially varying coefficient surface. Regions where lower and upper CI bounds exclude zero indicate statistically significant pathway relationships.

Surface comparison plots in the Supplementary **Figure S1** compared coefficient, lower CI, upper CI, and confidence interval width of surfaces, revealing regions of high uncertainty (wide confidence intervals) versus regions of stable, reproducible pathway relationships (narrow confidence intervals). These patterns suggest that certain anatomical compartments exhibit consistent neutrophil-lupus nephritis pathway crosstalk, while other regions show more variable or context-dependent relationships.

## 4. Discussion

Spatial transcriptomics has transformed our ability to study molecular processes within intact tissues, yet most analytical frameworks remain focused on identifying spatial domains, cell types, or individual genes [1]. Less attention has been given to modeling how genes and pathways interact across tissue space, despite the fact that disease processes and outcomes are often due to coordinated activity among multiple pathways rather than isolated molecular events. Towards this, we introduced the concept of a *spatial fingerprint* in this study.

Using a specified threshold (e.g., *Z*-score) to determine the statistically significant locations of a gene’s expression or the enrichment of a gene set (e.g., a cell type cluster, a curated pathway, a signature from an experiment) – or even a user-demarcated flexibly-shaped region – induces a complex 2D pattern defined as a spatial fingerprint on the tissue space. Given this generic definition, which could be based on genes or sets thereof, and the different logical and statistical operators that are applicable on a given fingerprint, recursively encoded fingerprints of increasing complexity could be systematically defined and analyzed by SFinx. Thus, information that is essentially visual and potentially subjective could now be transformed into a more objective representation. Indeed, a linguistic framework is being implemented on top of this new foundation in our ongoing work to connect with and leverage on the emerging AI models.

Here, we present SFinx along with SFDA, an integrated platform that further provides a continuous, functional representation of the discretely captured phenomenon. It reconstructs continuous pathway activity landscapes and quantifies spatially varying pathway-pathway and pathway-phenotype relationships. Application of SFinx to lupus nephritis highlighted the complex spatial organization of inflammatory signaling within diseased kidneys. Although neutrophil activation and lupus nephritis pathways exhibited areas of overlap, substantial portions of their activity remained spatially segregated, suggesting that inflammatory injury is organized into distinct molecular niches rather than a single homogeneous process. Spatial coefficient surfaces further demonstrated that associations between neutrophil activation and disease-related pathways vary considerably across tissue compartments, with both positive and negative relationships observed within the same section. These findings illustrate how spatially varying pathway interactions may contribute to localized tissue damage and underscore the importance of modeling disease mechanisms within their anatomical context.

A key technical advantage of this design is that aggregating genes into pathways before modeling should improve statistical stability compared to thousands of individual gene-wise tests. This is because rank-based gene-set scoring is inherently less sensitive to differences in sample composition, and penalized smoothers distribute information across neighboring tissue locations rather than treating each spot in isolation. These are principled expectations grounded in the properties of the underlying methods, not claims that have been formally benchmarked in the present study. A second advantage is that converting discrete spot measurements into continuous spatial surfaces makes cross-compartment and cross-section comparisons more tractable. Kriging achieves this in a principled way, returning both predicted values and associated uncertainty at every location rather than simply smoothing for visual appeal [30]. Third, and perhaps most importantly for translational applications, the outputs are inherently interpretable in biological terms. Pathway activity maps align directly with tissue architecture, overlap operators quantify whether two pathways occupy the same or distinct spatial domains, and spatially varying coefficient surfaces reveal not only the strength of association between two pathways but also where across the tissue that relationship shifts from positive to negative. This spatial resolution of pathway interactions is especially meaningful in disease, where the same signaling program can be adaptive in one compartment and pathogenic in another. The framework is broadly applicable to diseases characterized by spatially organized pathology. Potential applications include tumor microenvironment analysis in cancer, compartment-specific inflammation in autoimmune diseases, and lesion-centered molecular organization in neurodegenerative disorders [31–35]

Acknowledging the assumptions and limitations inherent in these analytical approaches is crucial. Deconvolution fundamentally remains an inverse problem that can become mathematically unstable when cell states are highly similar or when the assay is too sparse to support reliable de-mixing [14]. Regarding spatial modeling, while Kriging enhances spatial continuity for visualization, it cannot generate true biological resolution beyond the platform’s physical limits. It also risks over-smoothing sharp compartmental boundaries if the variogram is mis-specified or if inherently discontinuous biology is modeled as a continuous surface.

Similarly, Spatial Functional Data Analysis (SFDA) assumes that pathway effects vary smoothly across space. Consequently, the resulting confidence intervals should be interpreted as pointwise estimates rather than family-wise guarantees across the entire tissue surface [25,26]. Importantly, the derived coefficients quantify conditional spatial associations, not causality; they highlight regions where pathways co-vary in structured patterns, but do not establish mechanistic dependence between transcriptional pathways. The current implementation evaluates pairwise pathway relationships. Future extensions may incorporate higher-order interaction models capable of representing coordinated networks of multiple pathways simultaneously.

### 4.1. Future Directions

Future work will focus on extending SFinx beyond transcriptomic pathway mapping toward multimodal spatial inference. One promising direction is to obtain deeper insights into tissue phenotypes from histopathology images using pathway activity surfaces modeled with SFDA, enabling automated inference of molecular landscapes directly from routine tissue sections. In parallel, integrating SFinx with large language models (LLMs) and retrieval-augmented systems may facilitate interpretation of complex spatial pathway relationships by generating biologically grounded summaries and linking spatial findings to external biomedical knowledge. Together, these developments may help bridge spatial omics, computational pathology, and clinical decision support [31–33].

## 5. Conclusions

SFinx and SFDA provide a unified framework for transforming mixed-spot spatial transcriptomic measurements into continuous pathway landscapes and quantitative maps of spatial pathway interactions. By moving beyond spatially variable genes and tissue domains toward direct modeling of pathway-pathway and pathway-phenotype relationships, this approach enables a more mechanistic understanding of spatially organized disease processes. Application to lupus nephritis revealed anatomically localized inflammatory niches and compartment-specific pathway crosstalk, illustrating the biological insights that can emerge from pathway-centric spatial analysis. We anticipate that this framework will be broadly applicable across autoimmune diseases, cancer, neurodegeneration, and other conditions in which disease mechanisms are spatially heterogeneous and tissue-context dependent.

## Acknowledgments

Supported in part by the Intramural Research program at NIAMS/NIH (ZIA 041199). The contributions of the NIH authors were made as part of their official duties as NIH federal employees, are in compliance with agency policy requirements, and are considered Works of the United States Government. However, the findings and conclusions presented in this paper are those of the authors and do not necessarily reflect the views of the NIH or the U.S. Department of Health and Human Services.

---

**Supplementary Figure S1.**
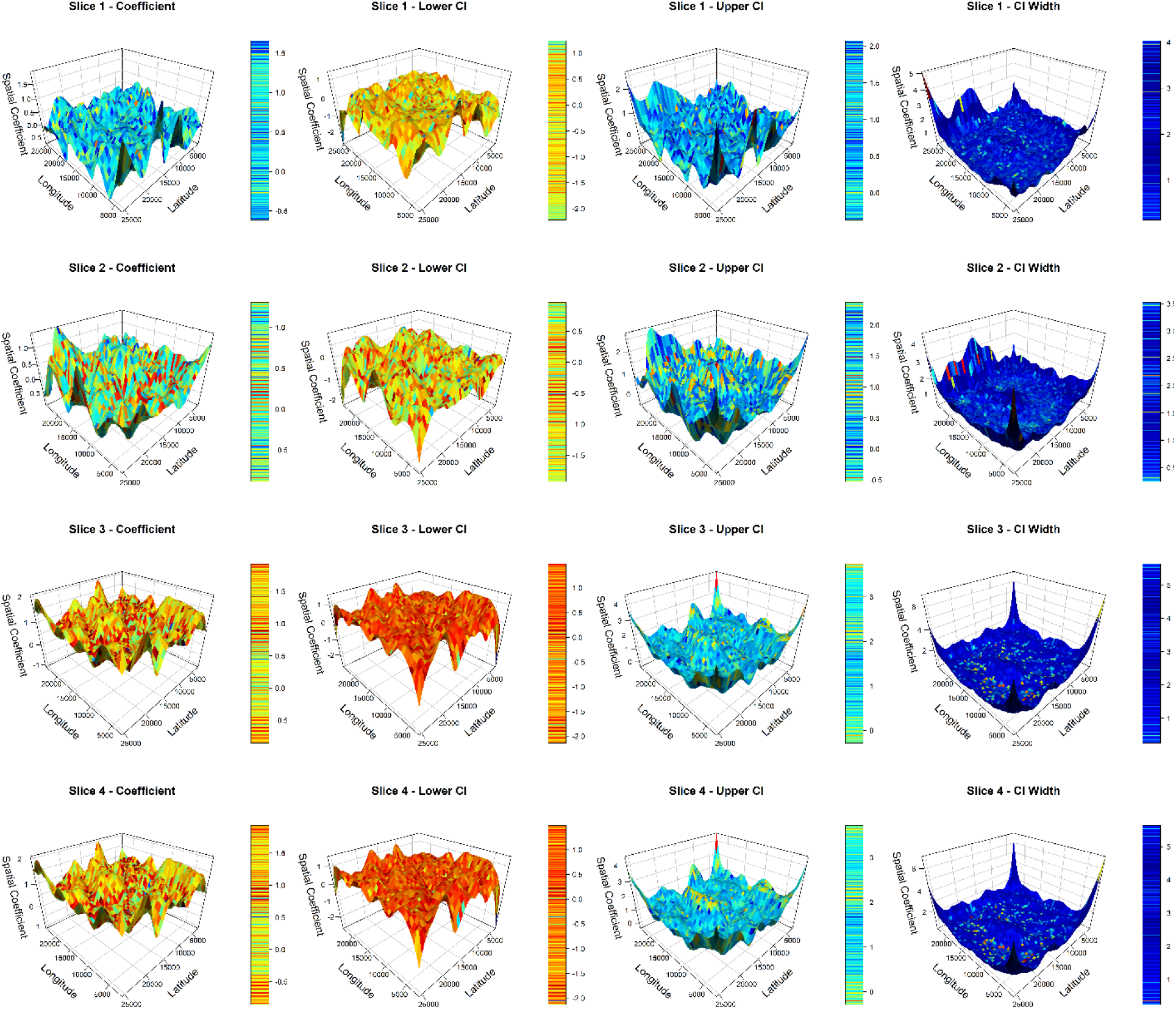
Evaluating Pathway Stability via Surface Coefficients. Wider intervals indicate regions of high uncertainty, while narrow intervals denote stable, reproducible pathway relationships.

